# The neural dynamics of novel scene imagery

**DOI:** 10.1101/429274

**Authors:** Daniel N. Barry, Gareth R. Barnes, Ian A. Clark, Eleanor A. Maguire

## Abstract

Retrieval of long-term episodic memories is characterised by synchronised neural activity between hippocampus and ventromedial prefrontal cortex (vmPFC), with additional evidence that vmPFC activity leads that of the hippocampus. It has been proposed that the mental generation of scene imagery is a crucial component of episodic memory processing. If this is the case, then a comparable interaction between the two brain regions should exist during the construction of novel scene imagery. To address this question, we leveraged the high temporal resolution of magnetoencephalography (MEG) to investigate the construction of novel mental imagery. We tasked male and female humans with imagining scenes and single isolated objects in response to one-word cues. We performed source level power, coherence and causality analyses to characterise the underlying inter-regional interactions. Both scene and object imagination resulted in theta power changes in the anterior hippocampus. However, higher theta coherence was observed between the hippocampus and vmPFC in the scene compared to the object condition. This inter-regional theta coherence also predicted whether or not imagined scenes were subsequently remembered. Dynamic causal modelling of this interaction revealed that vmPFC drove activity in hippocampus during novel scene construction. Additionally, theta power changes in the vmPFC preceded those observed in the hippocampus. These results constitute the first evidence in humans that episodic memory retrieval and scene imagination rely on similar vmPFC-hippocampus neural dynamics. Furthermore, they provide support for theories emphasising similarities between both cognitive processes, and perspectives that propose the vmPFC guides the construction of context-relevant representations in the hippocampus.

**Significance statement:** Episodic memory retrieval is characterised by a dialogue between hippocampus and ventromedial prefrontal cortex (vmPFC). It has been proposed that the mental generation of scene imagery is a crucial component of episodic memory processing. An ensuing prediction would be of a comparable interaction between the two brain regions during the construction of novel scene imagery. Here, we leveraged the high temporal resolution of magnetoencephalography (MEG), and combined it with a scene imagination task. We found that a hippocampal-vmPFC dialogue existed, and that it took the form of vmPFC driving the hippocampus. We conclude that episodic memory and scene imagination share fundamental neural dynamics, and the process of constructing vivid, spatially coherent, contextually appropriate scene imagery is strongly modulated by vmPFC.

## Introduction

Episodic memory formation and retrieval are long-established functions of the hippocampus (Scoville and Milner, 1957). However, cognitive impairments beyond recalling past experiences have been documented following hippocampal damage, including deficits in imagination and future-thinking (Hassabis et al., 2007a; Kwan et al., 2010; Kurczek et al., 2015). Accordingly, contemporary perspectives have converged on a more inclusive account of hippocampal function which accommodates the flexible construction of predictive or fictive representations (Hassabis and Maguire, 2007; Schacter and Addis, 2007; Eichenbaum and Fortin, 2009; Buckner, 2010; Maguire and Mullally, 2013).

One such interpretation, the Scene Construction Theory, proposes that the hippocampus constructs scene imagery to facilitate mental representations, whether recollected or imagined (Hassabis and Maguire, 2007; Maguire and Mullally, 2013). In this context, a scene is defined as a naturalistic three-dimensional spatially-coherent representation of the world typically populated by objects and viewed from an egocentric perspective (Maguire and Mullally, 2013; Dalton et al., 2018). In support of this thesis, functional MRI (fMRI) studies have revealed particularly anterior hippocampal recruitment while participants imagined novel scenes (Hassabis et al., 2007b; Zeidman et al., 2015; Zeidman and Maguire, 2016).

However, other regions, including the ventromedial prefrontal cortex (vmPFC), are recruited during (Hassabis et al., 2007b), and seem necessary for (Bertossi et al., 2016a), scene construction. An outstanding question, therefore, is how vmPFC interacts with hippocampus during the generation of scene imagery. The temporal resolution of magnetoencephalography (MEG) renders it a suitable method to address this question. Furthermore, a proposed mechanism of such inter-regional communication is oscillatory coherence (Fries, 2005). Increased theta synchrony between hippocampus and vmPFC has been observed during episodic memory retrieval (Fuentemilla et al., 2014) and integration (Backus et al., 2016), as well as memory-guided navigation (Kaplan et al., 2014) and decision-making (Guitart-Masip et al., 2013). Demonstrating analogous connectivity during the imagination of novel scenes would provide evidence that episodic memory and scene construction share not only similar loci of brain activity, but are supported by comparable network dynamics.

If oscillatory coherence between hippocampus and vmPFC is evident during scene construction, a question of further relevance concerns the direction of information flow between the two regions. Electrophysiological investigations in rodents have suggested that during initial contextual memory formation, hippocampal activation precedes that of vmPFC (Place et al., 2016). By contrast, retrieval (Place et al., 2016), detection of violations in learned information in humans (Garrido et al., 2015) and subsequent extinction, have been characterised by vmPFC driving hippocampus. It is unclear which pattern the generation of novel scene imagery might follow.

Campbell et al. (2018) investigated the directionality of information flow between hippocampus and vmPFC during the imagination of future events using dynamic causal modelling (DCM; Friston et al., 2003) of fMRI data. This revealed a greater influence of hippocampus over vmPFC. However, the temporal resolution of fMRI is not optimal to adequately characterise this dialogue. In contrast, McCormick et al. (2018) recently proposed that vmPFC initiates scene construction. From this perspective, the vmPFC would drive the hippocampus during the imagination of novel scenes.

The current study, we leveraged the high temporal resolution of MEG to address two questions. First, do the anterior hippocampus and vmPFC display coherent activity during the imagination of novel scenes relative to single objects? Given accumulating evidence that theta oscillations mediate the interaction between hippocampus and vmPFC, we predicted greater theta coherence between the two regions specifically for scene imagery. Second, does one of these regions exert a stronger influence over the other during scene imagination? Concordant with McCormick et al.’s (2018) proposal, we hypothesised that vmPFC would drive oscillatory activity in hippocampus. We asked participants to imagine novel scenes (and single isolated objects as a control condition) in response to single-word cues during MEG. A low-level baseline condition involving counting was also included. We then used a combination of source localisation techniques measuring power and coherence, as well as DCM for MEG to address the research questions.

## Materials and Methods

### Participants

Twenty-two participants (14 female) took part in this experiment (mean age 27 years; SD 7). Due to the verbal nature of the stimuli, only native English speakers were recruited. Participants gave written informed consent. The University College London Research Ethics Committee approved the study.

### Stimuli

Seventy-five scene words and 75 object words were used as stimuli for the imagination task. These comprised a sub-set of the stimuli devised by Clark et al. (2018) for a separate fMRI study. These word categories were closely matched on a number of properties (Table 1) to ensure any differences in neural activity could be solely attributed to the type of mental imagery they evoked. To enable vivid imagination, all words were rated as highly imageable (> 3.5/5). To facilitate the ease with which participants could construct the two different kinds of representations, words were designated as either scene or object-evoking if at least 70% of an independent sample of participants rated them as such (Clark et al., 2018). Object imagery was included as a suitable control condition for scene imagery, as vivid detailed mental imagery can be evoked and viewed from an egocentric perspective, without the requirement to construct a three-dimensional space. Furthermore, object imagery has been used as a closely matched control for scene imagery in previous neuroimaging studies (Clark et al., 2018; Hassabis et al., 2007b; Zeidman et al., 2015). Additionally, 75 number stimuli were also deployed in a third condition involving counting, which were matched to the scene and object words in terms of the number of letters and syllables. This condition served as a useful low-level baseline against which to compare the neural activity common to both scene and object imagery. Counting was preferred to a resting baseline, as such passive states have been associated with spontaneous neural activity in our regions of interest (Vincent et al., 2006).

**Table 1.**
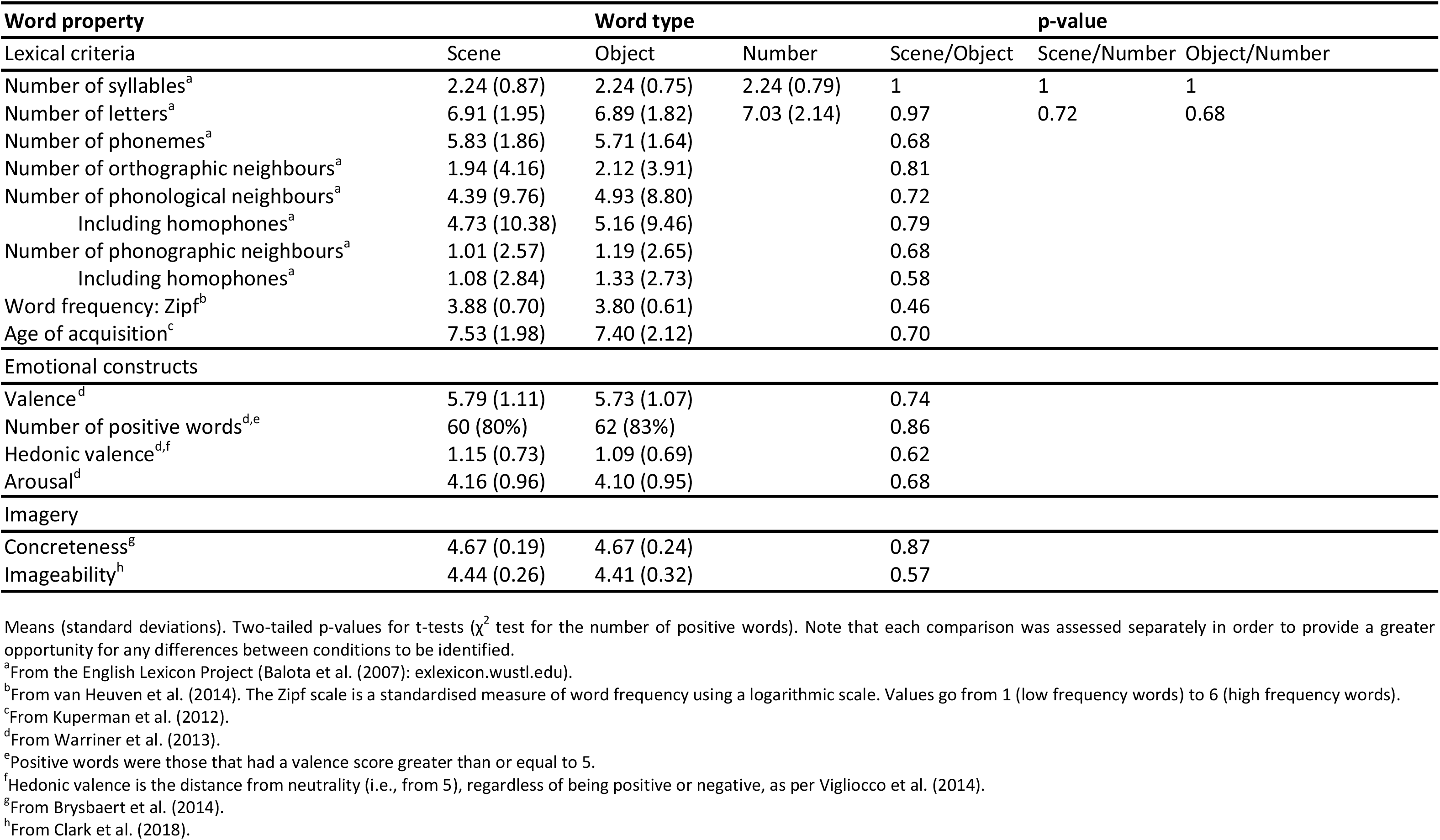
Properties of the word stimuli.

### Experimental design of the task

Prior to the MEG scan, participants received task instructions and practiced for the equivalent of two in-scanner sessions. For the scene imagination task, the instructions were as follows: “You will hear a word which evokes the mental image of a three dimensional space which you could step into, such as “jungle”. Then I want you to create as detailed a scene as you possibly can in your mind’s eye. I don’t want you to recall something from memory, such as when you visited such a place before. Instead, I want you to create the scene in your imagination. You will have just three seconds to do this, so please try and let it come as quickly and naturally as possible, and hold this image in your mind for the remainder of the three seconds”. For the object imagination task, participants were instructed to do the following upon presentation of an object word (e.g. “cushion”): “I want you to imagine a single object against a white background, as if it is floating in space. There should be nothing other than that object in your mind’s eye, so no background or anything associated with it; only the object. Also, try and make the object as large as possible, so that it takes up your entire field of view.” The counting baseline instructions were as follows: “count in threes from the presented number” (e.g. “forty”). In the MEG scanner, experimental stimuli were delivered aurally via MEG-compatible earbuds using the Cogent toolbox (www.vislab.ucl.ac.uk/cogent.php), running in MATLAB (version 2012).

To prepare them for each trial type, participants first heard either the word “scene”, “object” or “counting” (Figure 1). This was intended to minimise category confusion during the scene and object trials. Participants immediately closed their eyes and waited for an auditory cue which followed a jittered duration of between 1300 and 1700 milliseconds. As previously detailed in the instructions to participants, during scene trials participants constructed a novel, vivid scene from their imagination. During object imagery, participants imagined a single novel object, and counting trials involved mentally counting in threes from a number cue. The task periods were 3000 ms in duration. Participants then heard a beep and opened their eyes. They were presented with a rating screen. For scene and object trials, they were asked “What did you imagine?” If they failed to perform the task, they selected “unsuccessful”. Otherwise, they could select “low detail scene”, “high detail scene”, “low detail object” or “high detail object”. This allowed participants to indicate both the level of detail present in the mental imagery, and also to reclassify scene and object trials if they had inadvertently imagined an object as a scene or vice versa. For counting trials, participants were asked “How well did you concentrate?” on a scale from 1 (not at all) to 5 (extremely well). Following this was a 1000 ms delay before the next trial. There were eight scanner sessions in total, seven containing nine stimuli from each condition in a random order, and one final session with 12 stimuli from each condition. Most sessions contained 27 trials as this corresponded to the optimal time that participants could comfortably remain still and concentrate, which resulted in an excess of three trials per condition for the final session. Eighteen participants completed all eight sessions; the remaining four participants completed seven sessions due to technical issues with the recording equipment.

**Figure 1.**
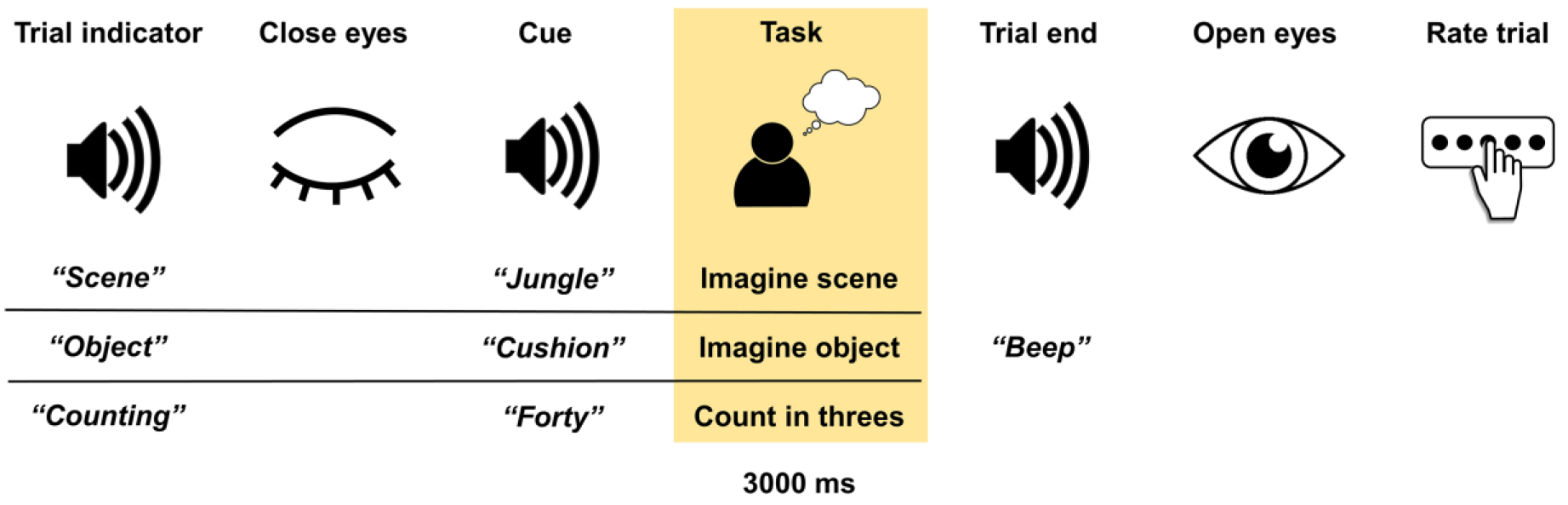
Trial structure. The task period selected for analysis is highlighted.

### Post-scan recognition memory test

After the scan, participants took part in a recognition memory test. Participants were not informed about this test prior to the scan to avoid confounding the imagination task with attempts to memorise the stimuli. They were presented with scene and object words, and asked if the word was previously presented in the scanner or not. The available response options were “yes” or “no”. Alongside the 75 scene and 75 object words which had been presented in the scanner, there were 38 scene and 38 object foils; the presentation was randomised for each participant. The scene and object foil words were also matched to the words presented in the scanner on the characteristics outlined in Table 1.

### Statistical analysis: behavioural data

#### Word properties

Comparisons between word category properties were performed using independent sample t-tests in the case of continuous variables and chi squared tests for categorical variables. Significance was determined at an alpha level of p < 0.05. Statistical analyses were performed using SPSS statistical package version 22 (SPSS, Chicago, IL).

#### In-scanner ratings

Comparison of the percentage of successful trials in each condition was performed using a repeated-measures one-way ANOVA. These percentages accommodate participants’ reclassifications of scenes and objects during task performance. Greenhouse–Geisser adjustment to the degrees of freedom was applied if Mauchly’s sphericity test detected a violation of sphericity. Comparison of trial reclassifications, and highly detailed ratings were performed using paired samples t-tests.

#### Post-scan recognition memory test

To ensure a relevant comparison with trials used in the MEG analysis, only successfully completed trials in the scanner were included as target stimuli in this analysis, and stimuli were reclassified as scenes or objects if the individual participant had imagined them as such during the task. Comparison of the percentage of correctly recognised scenes and objects, d’ and c (response bias) values were performed using paired samples t-tests.

### MEG recording and preprocessing

A CTF Omega whole-head MEG system with 273 functioning first order gradiometers recorded data at a sample rate of 1200 Hz. Four EOG electrodes were used to measure a participant’s vertical and horizontal eye movements. To rule out the possibility that differences between conditions in subsequent analyses were related to differences in eye movements, we computed the variance of these two EOG signals during each trial, which served as an indirect measure of saccadic activity. These variances were averaged across trials within each condition and normalised within subjects so that values for the three conditions summed to 1. A 1×3 repeated measures ANOVA did not detect any differences in eye movements between conditions (F_(2,42)_ = 1.12, *p* = 0.335). Data were epoched into 3 second mental imagery and counting periods, baseline corrected, and concatenated across sessions. For gamma (31-85 Hz) band analysis, a 50 Hz stop-band filter was applied to remove power line noise.

### Statistical analysis: MEG data

All MEG analyses were performed using SPM12 (www.fil.ion.ucl.ac.uk/spm). Source reconstruction was performed using the SPM DAiSS toolbox (https://github.com/spm/DAiSS).

### MEG source reconstruction

To estimate differences in power between experimental conditions in source space, the Linearly Constrained Minimum Variance (LCMV) beamformer was used. This filter uses a series of weights to linearly map MEG sensor data into source space to estimate power at a particular location, while attenuating activity from other sources. For each participant, a single set of filter weights was constructed based on the data from all three conditions within the 4-8 Hz band and a 0-3000 ms peri-stimulus window. Analysis was performed in MNI space using a 5 mm grid and coregistration was based on nasion, left and right preauricular fiducials. Co-registration and the forward model were computed using a single-shell head model (Nolte, 2003). Power was estimated in the theta (4-8 Hz) frequency band and the 0-3000 ms time window, with one power image per condition being generated for each participant. These images were smoothed using a 12 mm Gaussian kernel and entered into a second-level random effects (1×3) ANOVA in SPM to investigate power differences across conditions. Small volume correction was performed using a bilateral hippocampus mask generated from the AAL atlas implemented in the WFU PickAtlas software (http://fmri.wfubmc.edu/software/pickatlas). Identification of activation peaks in other regions was performed using the AAL atlas (Tzourio-Mazoyer et al., 2002). This analysis was repeated in the alpha (9-12 Hz) and gamma (31-85 Hz) bands to investigate task-based modulation of these frequencies.

We hypothesised that mental imagery would be associated with anterior rather than posterior hippocampal activation (Dalton et al., 2018; Hassabis et al., 2007b; Zeidman et al., 2015; Zeidman and Maguire, 2016). To test this hypothesis more thoroughly, we divided a left hippocampal mask into anterior, middle and posterior segments of equal length. We extracted the mean percentage difference in theta power between our imagery conditions and the counting baseline for each segment and participant. For each subject, we then performed a linear regression with hippocampal segment as the predictor variable and the difference in theta power from baseline as the dependent variable.

#### Coherence

Coherent activity between sources was measured using the Dynamic Imaging of Coherent Sources (DICS) approach (Gross et al., 2001). In this case, a beamformer reference signal estimate is first performed for a reference location, in this case a defined source in the anterior hippocampus. This analysis was performed separately in the 4-8 Hz and 9-12 Hz bands within the 3000 ms task time window. Then scanning the brain on a 3 mm grid, a signal estimate was made at each location and the coherence between this signal and the reference (in either the 4-8 Hz or 9-12 Hz bands) computed. These values were output as an image for each condition representing a brain-wide map of coherent activity with the reference source. These images were smoothed using a 12 mm Gaussian kernel, and contrasts between conditions at the group level were performed using a second-level random effects paired t-test in SPM.

For subsequent analyses, time series of theta activity during the 3000 ms imagery task period were extracted from two 10 mm radius spheres encompassing the hippocampal reference source and the group coherence peak in the vmPFC using the LCMV beamforming algorithm. To compare theta coherence between scene trials which were subsequently remembered in the post-scan recognition memory test, with scene trials that were forgotten, we performed a coherence analysis on the extracted time series of remembered and forgotten trials, via the *mscohere* function in the Matlab signal toolbox, using Welch’s averaged modified periodogram method. The imagination period was divided into three one second epochs with no overlap, and coherence was calculated over the frequency range of 4-8 Hz. One participant successfully recognised all 75 scene stimuli and was therefore excluded from this analysis. The same analysis was performed on scene trials which were imagined in high versus low detail. Given strong *a priori* hypotheses from previous research that theta coherence would be positively associated with subsequent memory performance (Backus et al., 2016), and highly detailed visual imagery (Fuentemilla et al., 2014), we performed a one-sided paired t-test in both cases.

#### Effective connectivity

To determine effective connectivity, we utilised DCM for Cross Spectral Densities (Moran et al., 2009), which analyses the magnitude of cross spectra between regions. The DCM approach involves creating a model specifying the direction of inter-regional information flow and fitting this model to the actual neural data. Multiple possible models can be generated and compared in the same manner to ascertain the best explanation for the experimental observations. DCM for MEG uses a biophysical neural-mass model which attempts to summarise the activity of millions of neurons within a region. This model accommodates different neuronal types and their intrinsic connectivity, treating the measurable output of these cell populations as a convolution of their input. If a region is being causally influenced by another, this activity should change in a predictable manner based on the nature of afferent input from the source. These inputs are characterised as forward, or “bottom up”, if they project to the middle layers of the cortex, backward, or “top down” if they target deep and superficial layers, or lateral if they innervate all layers (Felleman and Van Essen, 1991). One can therefore test biologically plausible models based on known structural connections between two regions, that differ in terms of which connections are functionally modulated by the experimental task.

The DCM estimation process attempts to fit these different models to the observed data as closely as possible by tuning their parameters. The evidence for any one model represents a balance between how accurate and parsimonious it is in explaining the data, as models with too many parameters are penalised. In this study, we used a convolution-based local field potential (LFP) neuronal model, as this is the simplest and most efficient approach when addressing hypotheses regarding differences in effective extrinsic connectivity (Moran et al., 2013). To assess which model best explains the observed data on a group level, random effects Bayesian model comparison (Stephan et al., 2009) is performed, which compares the evidence for each model across all participants, and generates the probability of it being the winning model. To assess the quality and consistency of model fit, we generated the log Bayes factor for each participant separately by computing the difference between the log evidence of the two models.

#### Event-related spectral perturbations

To further exploit the fine temporal resolution of MEG, we performed an additional analysis on the time-course of theta power changes across the scene imagination period. Using the same source-extracted time-series of activity in the vmPFC and anterior hippocampus, we applied a Morlet wavelet-based time frequency analysis with seven wavelet cycles across the 4-8 Hz frequency range. The time-period of interest extended from 500 ms prior to cue offset until the end of the 3000 ms imagination period (padded with real data). This was performed separately on each trial and subsequently averaged. The averaged time-frequency decomposition of the 0-3000 ms task period was then converted into log power, baseline corrected and transformed into dB values by rescaling to the 500 ms pre-task period. We then collapsed across frequency to produce a single time-series of event-related theta power changes in both regions. This time series was then smoothed using a 250 ms Gaussian kernel to attenuate variability in the temporal response across participants. The smoothed time series were entered into a random effects second level SPM analysis, generating a group F value at each time-point, with correction for multiple comparisons set at FWE p < 0.05.

## Results

### Behavioural

#### In-scanner task performance

Participants’ in-scanner self-rated performance was high (Table 2), with a minimal proportion of trials rated as unsuccessful. Of the trials eligible for subsequent analysis, there was a similar proportion of scene imagery, object imagery and counting trials (F_(1.22,25.64)_ = 2.25, *p* = 0.142). Scene and object stimuli appeared to successfully evoke the intended mental imagery, as a comparably low percentage of trials were reclassified in the two conditions (t_(21)_ = −1.67, *p* = 0.110). The majority of scenes and objects were imagined in high detail with both conditions matched on this rating (t_(21)_ = - 1.33, *p* = 0.199). Furthermore, for most counting trials, concentration was rated as high (>3/5), indicating participants successfully maintained their attention during this baseline condition.

**Table 2.** Scanner trial ratings (mean, SD).

| 922 |  | Condition |  |  |  | p-value |
| --- | --- | --- | --- | --- | --- | --- |
| 923 |  | Scene | Object | Counting | Unsuccessful | Scene/Object/Counting |
| 924 | Proportion of total trials (%) | 32.56 (2.51) | 31.03 (2.90) | 32.28 (1.24) | 4.13 (3.10) | 0.142 |
| 925 |  |  |  |  |  |  |
| 926 | Reclassified trials (%) | As Object | As Scene |  |  | Scene/Object |
| 927 |  | 3.03 (3.01) | 5.78 (8.15) |  |  | 0.110 |
| 928 |  |  |  |  |  |  |
| 929 |  |  |  |  |  | Scene/Object |
| 930 | High detail (%) | 71.41 (13.79) | 74.70 (12.71) |  |  | 0.199 |
| 931 |  |  |  |  |  |  |
| 932 | High concentration (%) |  |  | 78.09 (7.91) |  |  |

#### Post-scan recognition memory test

Participants were able to correctly recognise most of the scene and object stimuli that they previously imagined in the scanner, performing significantly above chance level in both conditions (Table 3). They remembered a greater proportion of objects than scenes (t_(21)_ = −2.10, p = 0.048). In addition, d’ scores indicated that participants were significantly less accurate in distinguishing between new and old scene compared to object words (t_(21)_ = −4.19, p < 0.001). Superior recognition memory for objects corroborates the findings of Clark et al. (2018), and implies that a greater power change or heightened connectivity during scene imagination relative to object imagination could not be attributable to better encoding of scenes. Of note, c values for both scene and object words indicated participants were conservative in their endorsement of recognised stimuli, in other words, they tended to only respond as such when they were confident in their response. Responses were more conservative for object than scene stimuli (t_(21)_ = −3.61, p = 0.002).

**Table 3.**
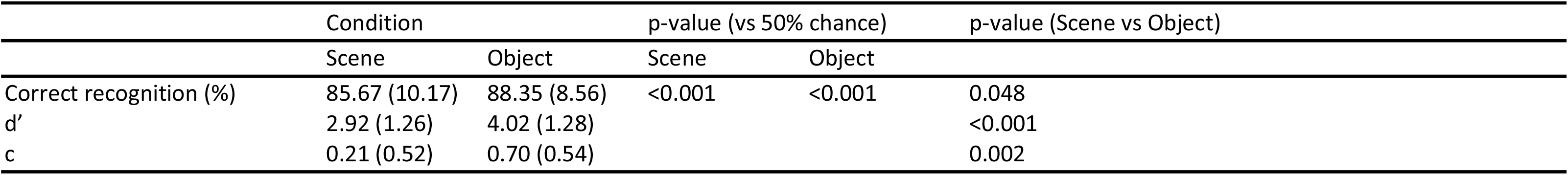
Post-scan recognition memory performance, d’ and c (response bias) values (mean, SD) for scenes and objects.

### MEG

#### Source space power changes during mental imagery

We first determined which brain regions were active during imagination in general, that is, scene and object imagination tasks combined compared to the low-level counting condition. Due to the obvious disparity in task demands between the imagery and baseline conditions, we used a conservative whole brain family-wise error corrected threshold of p < 0.001. A widespread change in theta power was observed during mental imagery when compared to the baseline task (Figure 2A). This was evident throughout the left anterior temporal lobe, with an activation peak at the whole-brain level in the inferior frontal gyrus (x = −38; y = 24, z = −4; Z-score = 6.23). Our primary *a priori* region of interest was the anterior hippocampus, where a significant change from baseline was also observed. A subsequent t-contrast revealed the observed changes represented an attenuation of theta power during imagination rather than an increase from baseline. Nonetheless, we regarded these power changes as an indication of task-related neural activity, and for subsequent connectivity analyses we did not exclude the possibility of inter-regional coherence in the presence of lower power. Subsequent small volume correction revealed an overall peak (−32, −4, −28; Z = 5.82) and sub-peak (−32, −6, −22; Z = 5.61) in the left anterior hippocampus.

**Figure 2.**
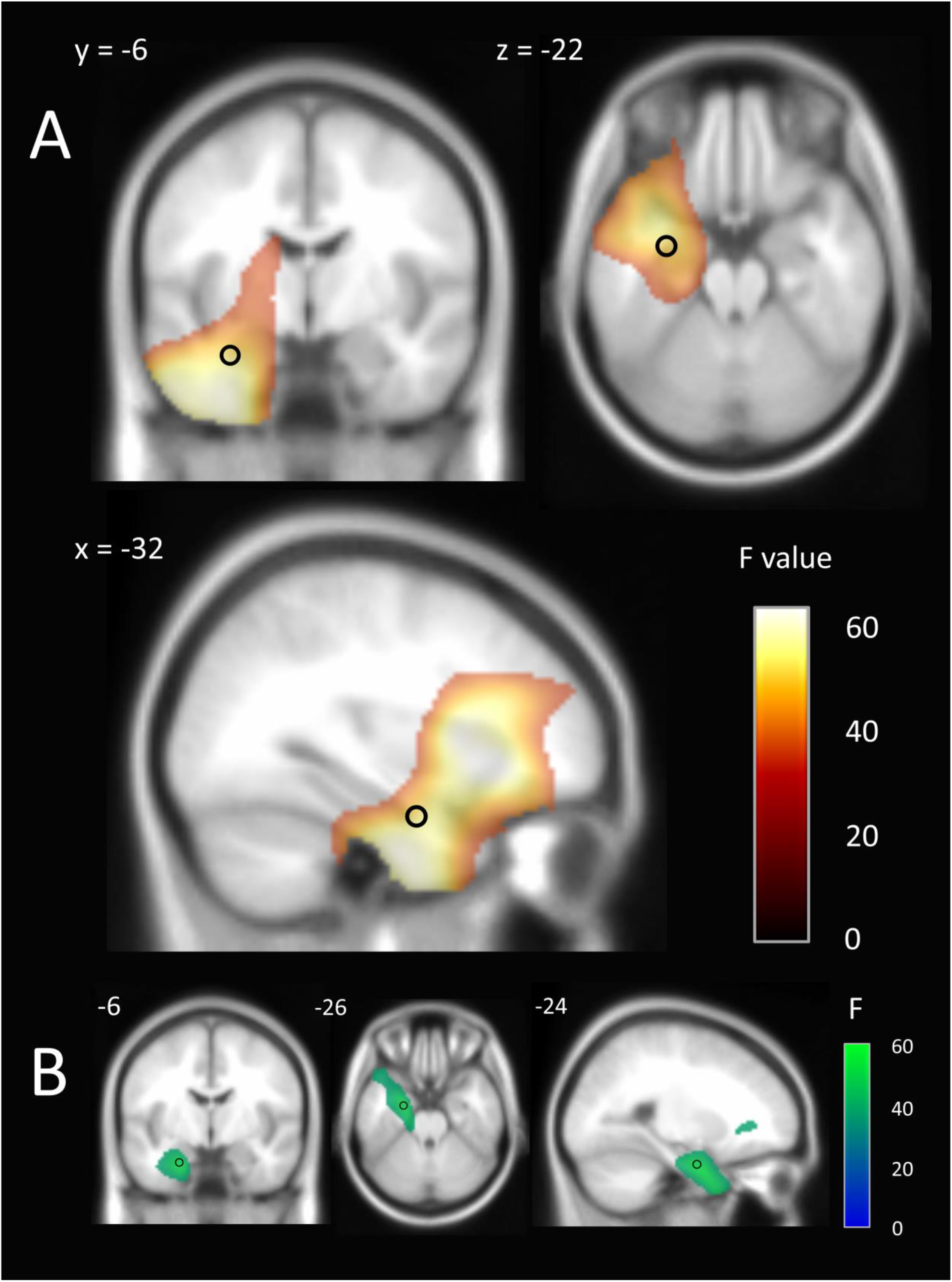
MEG source reconstruction of theta (4-8 Hz) and alpha (9-12 Hz) power changes during mental imagery (scenes and objects) compared to the baseline condition. The black circles represent the peak location of theta (A) and alpha (B) power changes in anterior hippocampus used for subsequent connectivity analyses. Images are FWE thresholded at p < 0.001 and superimposed on the Montreal Neurological Institute (MNI) 152 T1 image.

#### Gradient of theta power changes along the hippocampal longitudinal axis

Having observed changes in theta power in the anterior hippocampus at the group level, we investigated the consistency of this spatial selectivity across participants. We assessed each participant’s imagery-induced power change as a function of hippocampal segment (anterior, middle or posterior). Across the group, the regression was significant (F_(1,64)_ = 5.787, p = 0.019), with hippocampal segment explaining 8.3% of the variance in power difference between conditions. Fifteen of the 22 participants displayed a linear gradient of activation along the anterior to posterior hippocampal axis (Figure 3).

**Figure 3.**
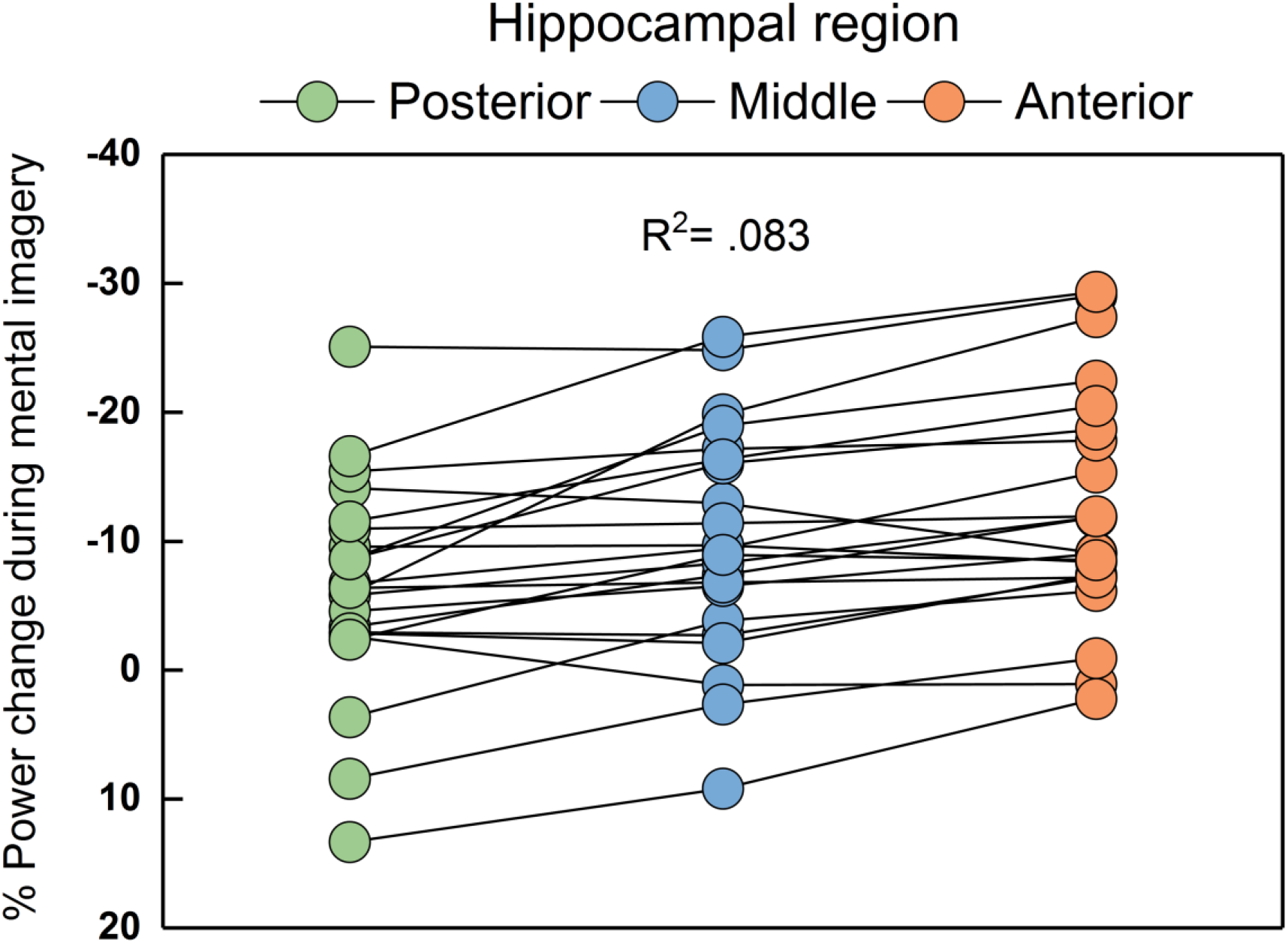
Gradient of theta power change along the hippocampal axis during mental imagery. The magnitude of theta power change increased significantly from posterior to anterior segments. This linear increase along the hippocampal axis was present in 15 of the 22 participants.

As the overall anterior hippocampal theta peak closely bordered the perirhinal cortex and fusiform gyrus, the more dorsal and posterior sub-peak (Figure 2A, black circle) was selected for connectivity analyses in order to be confident that source-localised activity originated from the hippocampus. This is because despite the fine spatial resolution of MEG beamforming, the volumetric full-width half-maximum (FWHM) of activation is on the order of at least a few millimetres (Barnes et al., 2004). Of note, no difference in theta power was observed at the whole brain level between scene and object imagination, even using an exploratory uncorrected threshold of p < 0.005 (cluster size > 5), or within the hippocampus in an ROI analysis (p < 0.05 uncorrected). In summary, scene and object imagery appeared to engage a common network of brain regions to a similar degree, including the anterior hippocampus. Therefore, any observed differences in connectivity between the two imagery conditions could not be explained by differences in power. Consequently, in the subsequent coherence analysis, we were able to directly explore the changes in network connectivity due to imagining scenes rather than single objects.

To investigate if there was a change in other frequency bands during our imagination tasks, we performed identical source localisation analyses across alpha (9-12 Hz) and gamma (31-85 Hz) frequencies. Significant changes in alpha power were observed in the medial temporal lobe, and small volume correction identified a single peak of activity in the anterior hippocampus (−24, −6, −26; Z = 5.53; Figure 2B). No differences in gamma power were observed between the imagery and baseline conditions at a FWE-corrected threshold of p < 0.05. The spatial proximity of alpha and theta power decreases in the anterior hippocampus allowed us to ascertain the specificity of the theta band rhythm in subsequent analyses of coherence, by using both peaks as a connectivity seed.

#### Hippocampal connectivity during scene imagery

Having established the peak locations of power changes in the anterior hippocampus during mental imagery, we then sought to investigate whether the imagination of scenes was associated with greater connectivity with any other regions of the brain when compared to object imagination. We found higher theta coherence in the left fusiform gyrus (Figure 4A; peak voxel: −32, −18, −32; Z = 3.58, p < 0.001 uncorrected) and parahippocampal cortex (peak voxel: −32, −38, −14; Z = 3.47, p < 0.001 uncorrected) during scene imagery. Given that we had a specific *a priori* hypothesis regarding connectivity between the hippocampus and vmPFC for scene imagery, we applied an uncorrected threshold of p < 0.005, and discovered a bilateral cluster of voxels coherent with the hippocampal source, at the most ventral extent of the vmPFC (Figure 4B; peak voxel: 18, 34, −16; Z = 2.87), with a sub-peak in the left vmPFC (−2, 46, − 28; Z = 2.86). This left-sided vmPFC peak was used for subsequent anatomically-informed analyses of effective connectivity as the hippocampus and vmPFC are connected ipsilaterally. Of note, the reverse contrast (greater connectivity for objects than scenes) did not reveal any significant results throughout the whole brain at a significance level of p < 0.005. Furthermore, at this threshold we did not observe higher coherence in the scene condition between the anterior hippocampus and other regions previously implicated in scene construction such as the precuneus, retrosplenial cortex, calcarine sulcus or occipital gyrus.

**Figure 4.**
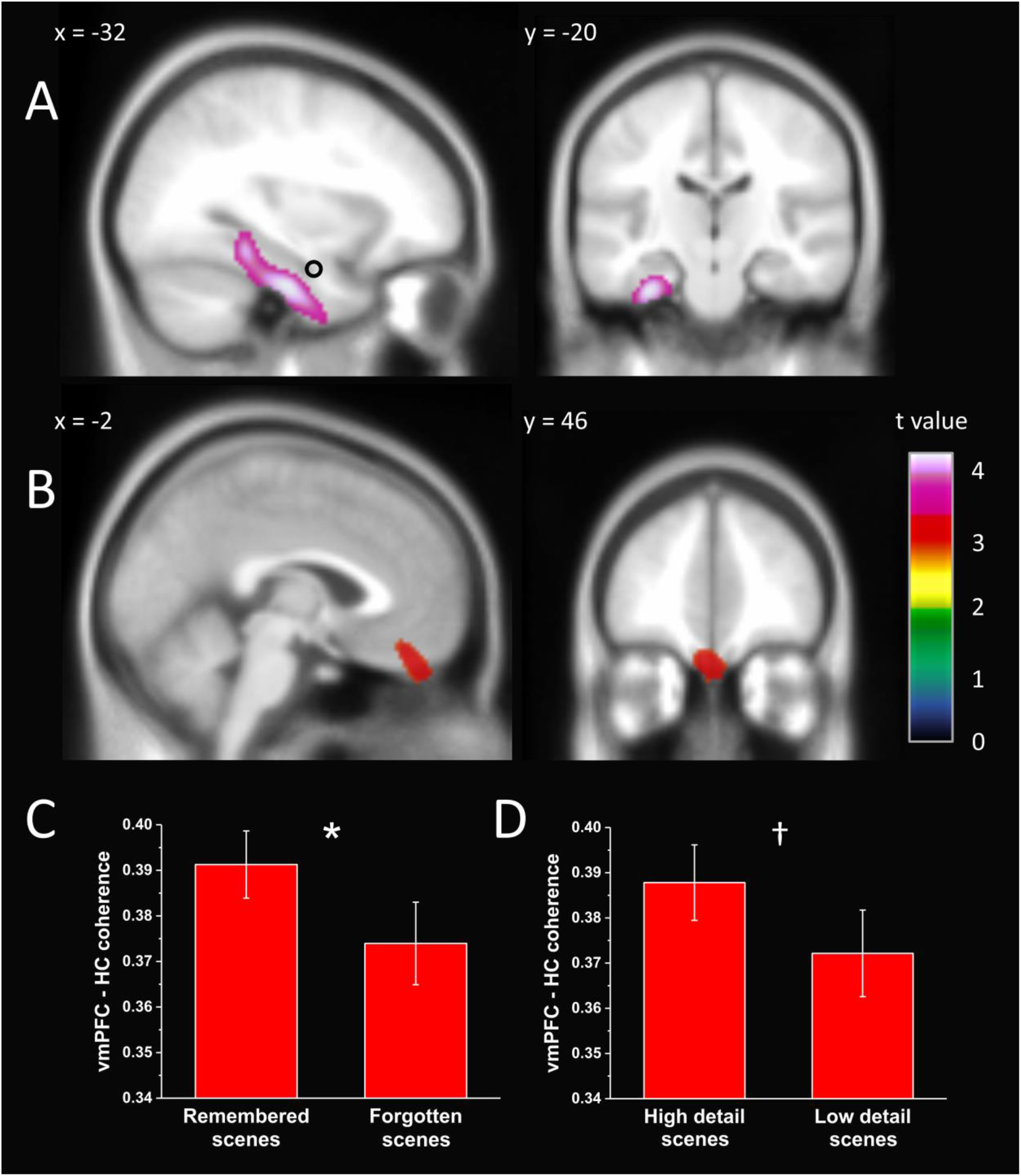
Brain areas displaying higher theta coherence with the left anterior hippocampus during scene imagination compared to object imagination. ***A***, The fusiform and parahippocampal cortices showed higher coherence with the hippocampal source (black circle), display thresholded at p < 0.001 uncorrected, superimposed on the MNI 152 T1 image. ***B***, The vmPFC, display thresholded at p < 0.005 uncorrected, also showed higher coherence with the hippocampal source (black circle). No areas showed higher theta coherence with the hippocampus for object over scene imagery. ***C***, Coherence between the vmPFC and hippocampus was significantly higher for subsequently remembered than forgotten scenes, *p = 0.018. ***D***, A similar trend was observed for scenes imagined in high versus low detail, †p = 0.058. Error bars represent ± 1 SEM.

To determine whether this increased coherence for scenes over objects was specific to the theta oscillation, we also used the location of peak alpha power change in the anterior hippocampus as a seed for brain-wide coherence in the alpha band. We did not find any region coherent with the hippocampus at a threshold of p < 0.001 (uncorrected), nor was coherent activity with the vmPFC observed at a threshold of p < 0.005 (uncorrected).

Our primary hypothesis was that scene construction and episodic memory are subserved by similar interactions between the hippocampus and the vmPFC. To make this direct comparison, we compared the theta coherence of imagined scenes which were subsequently remembered in the post-scan recognition test, with those that were forgotten. In support of our hypothesis, coherence between the two regions was higher for imagined scenes which were also successfully memorised (t_(20)_ = 2.25, p = 0.018; Figure 4C), despite the absence of any explicit instruction to do so. One potential interpretation of higher coherence in the scene construction condition is that it is a more effortful task than object construction. If this were the case, scenes imagined in low detail, which arguably serves as a proxy for difficulty, would show greater connectivity between the hippocampus and vmPFC. However, the opposite trend was observed, where coherence between the two regions was greater for high detail when compared to low detail scenes, a difference which approached significance (t_(21)_ = 1.64, p = 0.058; Figure 4D).

#### Effective connectivity during scene imagery

Having established higher theta coherence between hippocampus and vmPFC during scene imagination, we then investigated the directionality of information flow between the two regions. Using DCM for Cross Spectral Densities, we first specified a biologically plausible model of hippocampal-vmPFC connectivity. As the anterior hippocampus projects directly to the middle layers of the ventral extent of the ipsilateral vmPFC (where we observed high theta coherence) via the fornix (Aggleton et al., 2015), we designated this connection as forward. Because return projections are indirectly channeled through the entorhinal cortex (EC), vmPFC influence over the hippocampus is best characterised by the pattern of laminar innervation in EC. vmPFC efferents terminate in all layers of EC (Rempel-Clower and Barbas, 2000), and consequently we designated the return connection as lateral (Felleman and Van Essen, 1991). Additional thalamic relays exist between the two structures (Varela et al., 2014; Xiao et al., 2009) but are not considered here as the hierarchical nature of this connection type is unclear (Bastos et al., 2012).

Of key interest was the predominant direction of information flow between hippocampus and vmPFC during scene imagination. Therfore we proposed two anatomically informed models. In model 1, hippocampal activity drove the vmPFC via its forward connection. In model 2, lateral projections from the vmPFC modulated activity of the hippocampus (Figure 5A). The specified time period was the 3000 ms imagination task, and analysis of cross spectral density was constrained to the theta (4-8Hz) frequency band to retain consistency with previous power and coherence analyses. We applied both models to the observed data, and subsequently performed Bayesian model comparison to determine which model was most likely to explain the relationship between the two regions. The model most likely to be the winning model across all subjects, with a probability of 97.66%, was the vmPFC exerting a causal influence over the anterior hippocampus during the imagination of novel scenes (Figure 5B).

**Figure 5.**
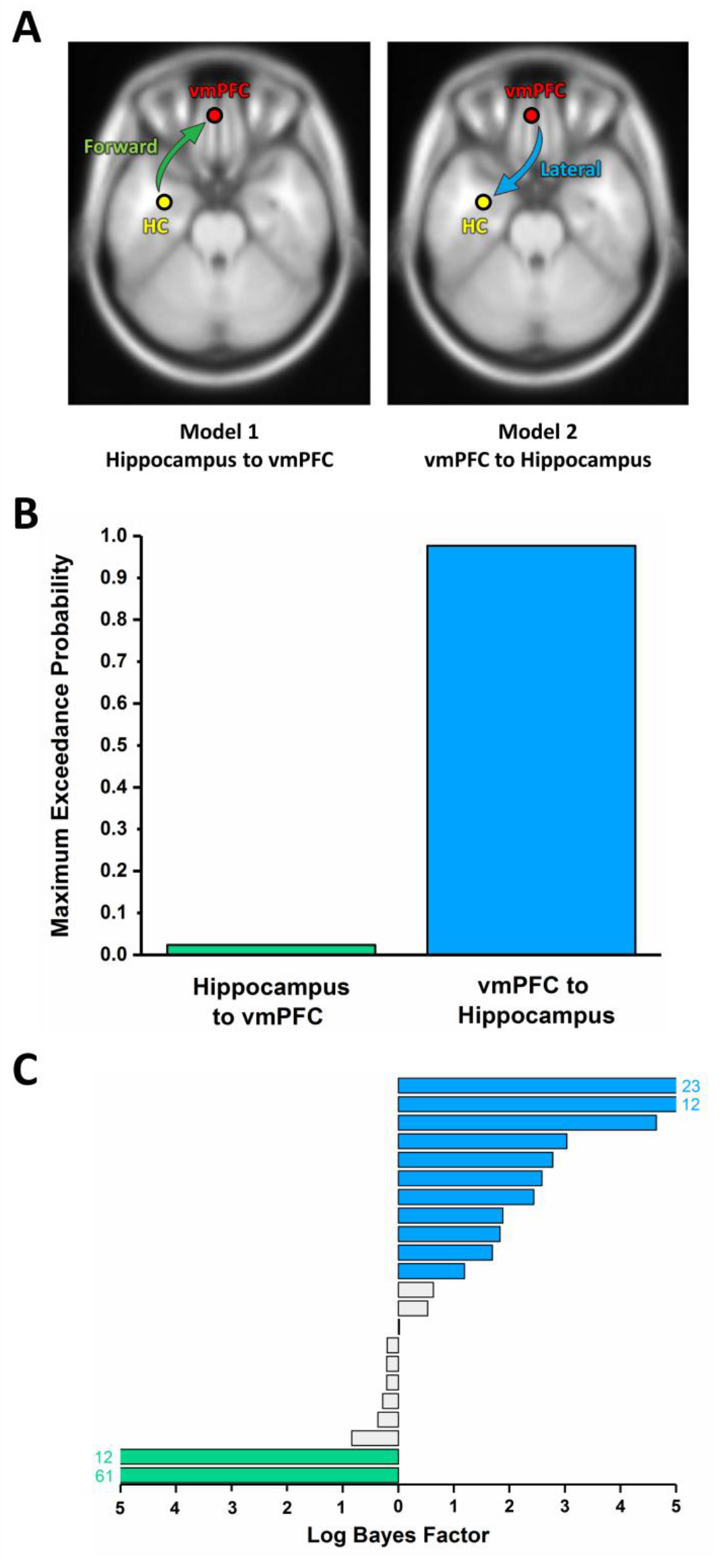
Dynamic Causal Modelling of the interaction between the hippocampus and vmPFC. ***A***, Two proposed models of effective connectivity between the coherent peaks in hippocampus and vmPFC. ***B***, Results of Bayesian Model Comparison indicated a stronger influence of the vmPFC on hippocampal activity during scene imagination. ***C***, Log Bayes factors for each participant. Blue bars indicate positive to strong evidence for vmPFC driving hippocampus, the model which most consistently fit across participants. Green bars represent the only two cases where evidence of the hippocampus driving vmPFC was observed. Grey bars represent the remaining participants where there was no conclusive evidence for either model. Where log Bayes factors exceeded five, bars are truncated and the exact values are adjacently displayed.

To quantify the consistency of model fit, we calculated the log Bayes factor for each model and participant separately (Figure 5C). According to the classification of Kass and Raftery (1995), a Bayes factor of 3 to 20 (log equivalent 1.1 to 3) constitutes positive evidence in favour of a model, with higher values indicating strong evidence. Eleven of the 22 participants in this study displayed positive or strong evidence for vmPFC driving hippocampus (Figure 5C, blue bars). In contrast, evidence for hippocampus driving the vmPFC was only present in two participants (green bars). In the remainder of the sample, there was no conclusive evidence for either model (grey bars).

#### Event-related spectral perturbations

In a complementary analysis to fully leverage the temporal precision of MEG, we compared the timing of neural responses in the vmPFC and hippocampus during the scene construction task. We charted the change in theta power over three seconds of scene imagery, relative to 500 ms preceding the offset of the cue (Figure 6). Overall peak activations (at a FWE-corrected threshold of p < 0.05) of the hippocampus (806 ms, Z = 3.72, bold yellow line) and vmPFC (899 ms, Z = 3.77, bold red line) were temporally proximal. However a much earlier sub-peak was observed in the vmPFC 310 milliseconds (Z = 3.19) following the offset of the cue. An additional peak of activity was observed in the vmPFC at a later point in the trial (1752 ms, Z = 3.68). The rapid initial engagement of the vmPFC in contrast to the slower kinetics of hippocampal activation provides additional evidence that the vmPFC may drive hippocampal activity to facilitate the construction of scene imagery.

**Figure 6.**
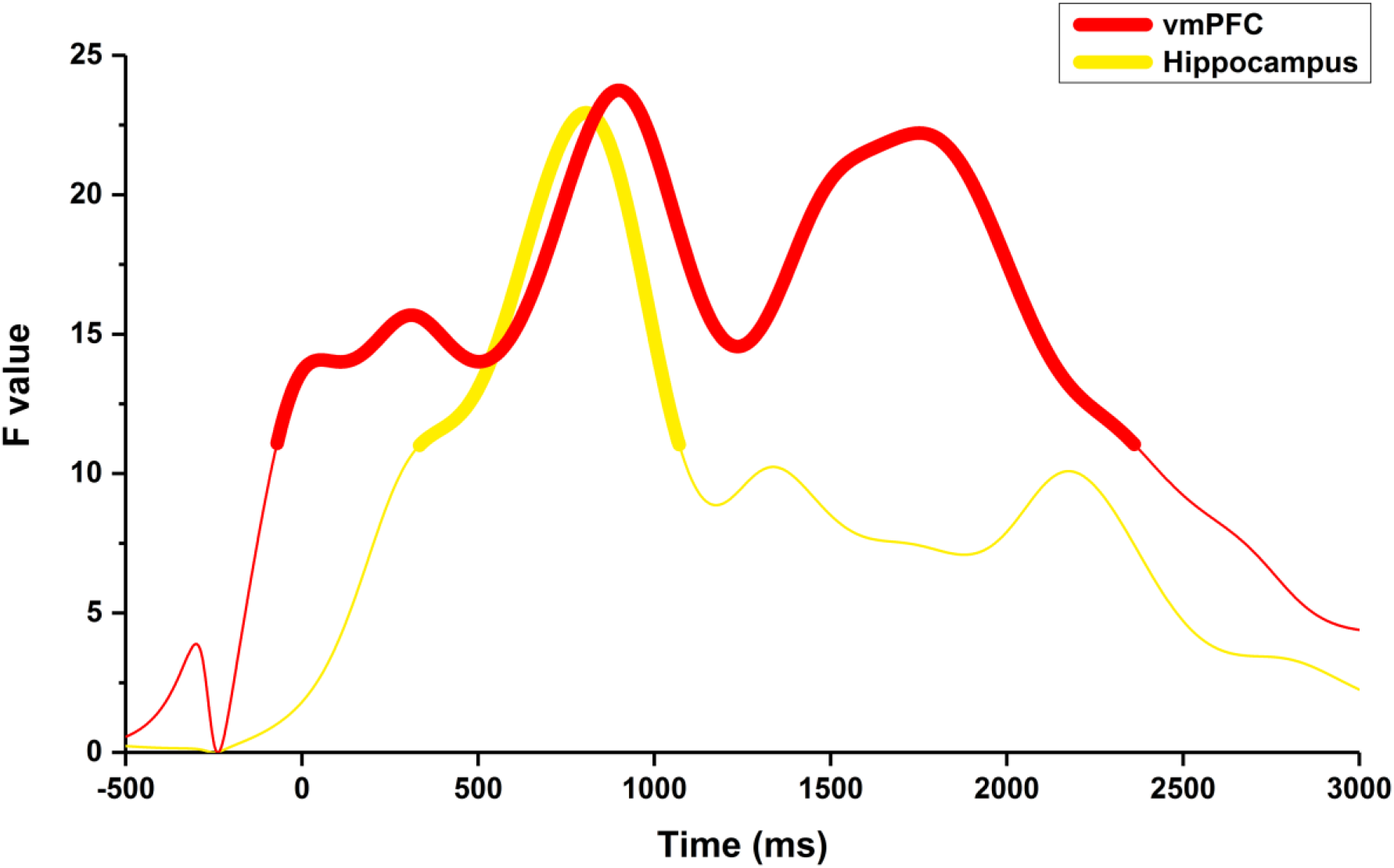
Temporal profile of theta power changes in the vmPFC and hippocampus during scene construction. Line segments in bold indicate periods of significant (p < 0.05 FWE corrected) changes in power relative to the pre-imagination period for both the vmPFC (red) and hippocampus (yellow). An initial peak in vmPFC activity was observed 310 milliseconds following cue offset, consistent with a role for the vmPFC in driving hippocampal activity. Maximal activation of both regions occurred within a narrow temporal window (hippocampus: 806 ms, vmPFC: 899 ms). A peak of similar magnitude was evident in the vmPFC at a later stage in the imagination period (1752 ms).

## Discussion

This study investigated whether the imagination of novel scenes is supported by a hippocampal-vmPFC dialogue. Mental imagery, whether of spatially-coherent scenes or isolated objects, resulted in comparable theta power decreases in the left anterior hippocampus. However, theta coherence between anterior hippocampus and vmPFC was significantly higher for scene compared to object imagination, and was of a greater magnitude for scenes that were subsequently remembered, or imagined in high detail. The observed coherence was also specific to the theta band. In addition, DCM of this interaction revealed the vmPFC drove hippocampal activity during the scene construction process, and an analysis of event-related spectral perturbations showed that the activity of vmPFC preceded that of hippocampus.

These findings corroborate fMRI studies demonstrating that the anterior hippocampus contributes to the mental construction of novel scene imagery (Hassabis et al., 2007b; Zeidman et al., 2015; Zeidman and Maguire, 2016, Dalton et al., 2018). An interesting feature of our data, which contradicts some previous reports of theta increases during learning and episodic memory tasks, is the attenuation of this frequency during imagination. However, evidence has accumulated from electroencephalography (EEG; Fellner et al., 2016) and MEG (Guderian et al., 2009) demonstrating a strong decrease in medial temporal lobe theta during episodic memory encoding. These findings have been validated using direct intracranial recordings in humans, with brain-wide decreases in theta power predicting subsequent recall (Burke et al., 2013; Greenberg et al., 2015), and specifically in the hippocampus (Sederberg et al., 2007; Lega et al., 2012; Matsumoto et al., 2013; Lega et al., 2017). A decrease in 8Hz power has also been reported during episodic memory retrieval (Michelmann et al., 2016). Furthermore, decreases in low frequency power appear to be negatively correlated with the fMRI BOLD response (Fellner et al., 2016; Scheeringa et al., 2011), and, therefore, our findings may be consistent with observed BOLD increases in previous fMRI studies.

The functional significance of the observed theta power decrease is not yet clear. However, in rodents, reduced hippocampal theta power is observed upon introduction to a novel or unexpected environment (Jeewajee et al., 2008). Our task involved the rapid mental construction of novel scenes (and so environments) in response to unpredictable stimuli, and the underlying oscillatory dynamics may, therefore, be similar. Importantly, a lower amplitude signal can still contain rich information about the underlying mental representations during episodic memory retrieval (Michelmann et al., 2016).

In the current study, theta power in the hippocampus did not differentiate between scene and object imagination. This accords with recent fMRI findings showing that object stimuli engage the hippocampus (Clark et al., 2018; Dalton et al., 2018), and that different task and stimulus-specific circuits may exist within the hippocampus, which receive distinct cortical afferents. Concordantly, Fuentemilla et al. (2014) demonstrated that while hippocampal theta power was similar during the retrieval of semantic and autobiographical memories, connectivity with the vmPFC was higher during the latter hippocampal-dependent task. Likewise, we observed increased theta coherence between the anterior hippocampus and vmPFC during scene, more so than object, construction, indicating a similar network dynamic may support episodic memory retrieval and scene imagination. Providing further support for such a shared dynamic, we discovered that the magnitude of coherence between these two regions during scene imagination predicted whether or not the stimulus was subsequently remembered.

How might the vmPFC facilitate both processes? The vmPFC is a proposed target of systems-level memory consolidation and long-term storage (Nieuwenhuis and Takashima, 2011), with evidence in humans of the reactivation of specific remote autobiographical memory traces (Bonnici et al., 2012; Barry et al., 2018). Yet participants in the current study were instructed to avoid recalling a specific autobiographical memory during the imagination process. However, the vmPFC is also thought to slowly extract regularities across past experiences to form superordinate representations, or schemas (van Kesteren et al., 2013; Gilboa and Marlatte, 2017). These may serve as flexible conceptual “templates” within which to rapidly and efficiently construct spatially coherent novel scenes in concert with the hippocampus.

Patients with vmPFC damage have schema-related deficits (Ciaramelli et al. 2006; Gilboa et al, 2006; Ghosh et al., 2014; Warren et al., 2014) and are impaired at constructing scene imagery (Bertossi et al., 2016a; Bertossi et al., 2016b; Bertossi et al., 2017; De Luca et al., 2018). However, such patients can generate imagery for individual scenes from autobiographical events in response to highly specific cues (Kurczek et al., 2015). Consequently, and in keeping with its role in supporting schema, it has been suggested that the vmPFC is necessary to select appropriate elements for a particular scene, while the hippocampus is needed to construct the scene imagery (McCormick et al., 2018). Our results are compatible with this interpretation, as coherence between the two regions was of a greater magnitude for scenes which were imagined in high detail.

An ensuing question is how the two brain regions collaborate to produce these integrated representations. One interpretation is that during imagination, the role of the vmPFC is to fuse distributed knowledge into a novel representation (Benoit et al., 2014), with corresponding evidence from fMRI that the hippocampus drives activity in the vmPFC when simulating the future (Campbell et al., 2018). However, an alternate perspective, as alluded to above, holds that the vmPFC exerts direct control over the hippocampus to select context-relevant representations (Eichenbaum, 2017; McCormick et al., 2018). Our finding of vmPFC exerting a causal influence over the hippocampus during scene imagination is more consistent with this latter view. We also regard MEG as a method well-suited to characterising this relationship given that it is a direct and time-resolved measure of neural activity.

What aspects of the lateral projection from vmPFC are influencing the hippocampus remains an open question. The preferential termination of neurons in the middle layers of the entorhinal cortex (Rempel-Clower and Barbas, 2000) indicates the presence of a driving and excitatory input (Bastos et al., 2012) to the hippocampus. Conversely, the majority of residual connections project to inhibitory neurons in the deepest layers (Joyce and Barbas, 2018), suggesting vmPFC also heavily constrains hippocampal output. From a behavioural perspective, the inability of patients with vmPFC damage to retrieve context-relevant information during scene construction tasks (Bertossi et al., 2016a) – which involve cues that are relatively unconstrained (Hassabis et al., 2007a) – while being also unable to suppress context-irrelevant information while confabulating (Turner et al., 2008), suggests the vmPFC may control both hippocampal input and output during scene imagination. Our time-resolved analysis of vmPFC activity during the construction of novel scenes identified additional periods of early and late engagement relative to hippocampal activation, adding credence to this view.

Our results also revealed that engaging in mental imagery in response to scene and object words relative to the counting baseline resulted in power decreases in the left inferior frontal gyrus. This activation peak was localised to Brodmann Area 47, a region implicated in the processing of verbal stimuli (Krieger-Redwood et al., 2015), in particular single words (Cutting et al., 2006). As this activation was common to scene and object words, it likely reflects the increased demands in semantic processing relative to the number stimuli in the baseline counting condition. This lexical processing is likely to be contemporaneous with mental imagery (Lewis and Poeppel, 2014), and it is therefore unlikely that word comprehension and imagination in the current experiment are temporally dissociable.

One additional finding which differentiated scene from object construction was increased theta coherence between the anterior hippocampus and the fusiform and parahippocampal cortices. Increased fusiform activity has also been observed in a separate fMRI study involving the scene word stimuli used in the current experiment (Clark et al., 2018). This region appears to represent diverse categories of objects, living beings and their interactions (Grill-Spector et al., 2006; Cukur et al., 2013), and scenes represent the coherent integration of these constituent elements. Observed coherence with the parahippocampal cortex during scene imagination is consistent with the parahippocampal cortex’s proposed role in processing scenes (Epstein, 2008; Mullally and Maguire, 2011). This connectivity profile, therefore, suggests that the anterior hippocampus may be a convergence zone for appropriate object categories and their interaction within a defined space, permitting the generation of scene imagery (Dalton and Maguire, 2017).

In summary, our results characterise, for the first time, a core neural dynamic which underlies scene construction, a process held by some to be fundamental to key cognitive functions including episodic memory and future-thinking. Previous studies have demonstrated co-activation of (Hassabis et al., 2007b), and a dependency on (Hassabis et al., 2007a; Bertossi et al., 2016a), the hippocampus and vmPFC during scene construction. By leveraging the high temporal resolution of MEG, we have extended these findings to demonstrate their functional connectivity during this process. Furthermore, we have shown that the direction of information flow during scene imagination mirrors that observed during episodic memory retrieval (Place et al., 2016), with vmPFC driving hippocampal activity. We conclude that episodic memory and imagination share fundamental neural dynamics, and the process of constructing vivid, spatially coherent, contextually appropriate scene imagery is strongly modulated by the vmPFC.

## Acknowledgements

This work was supported by a Wellcome Principal Research Fellowship to E.A.M. (210567/Z/18/Z) and the Centre by a Centre Award from Wellcome (203147/Z/16/Z). The authors are grateful to David Bradbury for technical assistance.

